# KILDA: identifying KIV-2 repeats from kmers

**DOI:** 10.1101/2025.01.17.631891

**Authors:** Molitor Corentin, Labidi Timothy, Rimbert Antoine, Cariou Bertrand, Di Filippo Mathilde, Bardel Claire

## Abstract

**Motivation:** High concentration of lipoprotein(a), a lipoprotein with proatherogenic properties, is an important risk factor for cardiovascular disease. This concentration is mostly genetically determined by a complex interplay between the number of Kringle-IV type 2 repeats and Lipoprotein(a)-affecting variants. Besides lipoprotein(a) plasma concentration, there is an unmet need to identify individuals most at risk based on their *LPA* genotype.

**Results:** We developed KILDA, a Nextflow pipeline, to identify the number of Kringle-IV type 2 repeats and Lp(a)-affecting variants directly from kmers generated from FASTQ files. The pipeline was tested on the 1000 Genomes Project (n=2459) and results were equivalent to DRAGEN-LPA (R^2^=0.93). *In-silico* datasets proved the robustness of KILDA’s predictions under different scenarios of sequencing coverage and quality.

**Conclusion:** KILDA is an open-source and free-to-use pipeline to identify the number of Kringle-IV type 2 repeats and lipoprotein(a)-associated variants. Its results are equivalent to DRAGEN-LPA, offering a free and robust tool for determining the *LPA* kringle number even when inputting low coverage libraries.

**Availability:** KILDA is publicly available at https://github.com/HCL-HUBL/KILDA along with a recipe to build an Apptainer image containing all the required dependencies.

**Supplementary information:** Supplementary data are available at *Bioinformatics* online.

## 1 Introduction

Cardiovascular disease (CVD) is one of the leading causes of death worldwide and is on the rise [1]. Concentration of lipoprotein(a) (Lp(a)) in the blood is an independent and important risk-factor for CVD [2], [3]. Lp(a) is composed of a low-density lipoprotein-like (LDL-like) particle in which apoB is bound to apolipoprotein(a) (apo[a]), the pathognomonic component of Lp(a). The Lp(a) concentration remains stable across the lifespan of an individual and is mostly genetically determined, by a complex interplay between genetic variants and the number of repeats of Kringle IV type 2 (KIV-2) [4]. KIV-2 is a repeated region, ∼5.5 kbp long, with a highly variable number of copies between individuals. Importantly, the number of KIV-2 repeats is inversely correlated with Lp(a) plasma levels [5]. Despite recommendations from the European and French societies [3] [6] Lp(a) measurement is not performed routinely in clinical practice, neither systematically nor in subjects at high cardiovascular risk. This is mainly due to the fact that medical laboratories are not equipped to measure Lp(a) and that such analyses are costly and not necessarily reimbursed.

Moreover, the diagnosis of familial hypercholesterolemia (FH) is characterized by genetically defined high levels of LDL-cholesterol. LDL-cholesterol plasma levels are, in most cases, calculated from formulas using total cholesterol [7]. The cholesterol content of Lp(a) can interfere with these equations [8]. Consequently, individuals with elevated Lp(a) levels may be erroneously diagnosed with FH and could lead to unnecessary FH genetic testing [9]. When these tests yield negative results, the subsequent cascade screening, a process to identify affected family members, is not initiated. This oversight, in light of coming Lp(a)-lowering drugs [10], may result in missed opportunities for early detection and intervention in truly affected individuals and their relatives.

An open question remains: How can we efficiently detect, using a genetic approach, hypercholesterolemic patients most likely to have elevated Lp(a) concentration? As Lp(a) concentrations are highly heritable, leveraging latent genetic information provide a promising approach to pinpoint these patients. For this purpose, the *LPA* gene and variants that can predict circulating concentrations of Lp(a) [11] must be sequenced and a tool is required to accurately estimate the KIV-2 copy numbers. Recently, the DRAGEN-LPA caller [12] managed to get accurate estimation of KIV-2 copy numbers. However, DRAGEN-LPA has been developed to process whole genome sequencing data, which is not ideal to use in molecular diagnosis, due to costs and closed-source code limitations.

Here, we present KILDA (KIv2 Length Determined from a kmer Analysis), an open-source and freely available pipeline, written in Nextflow and python. KILDA is designed to estimate KIV-2 copy numbers and detect the presence of Lp(a)-affecting variants directly from FASTQ files. We benchmarked our tool against DRAGEN-LPA using the 1000 Genomes Project dataset. Additionally, we evaluated KILDA’s accuracy using simulated samples with predetermined KIV-2 repeat counts, testing its robustness under various constraints of sequencing coverage and quality.

## 2 Methods

KILDA estimates KIV-2 repeat numbers by analysing the ratio of kmer occurrences between the KIV-2 region and one or more normalisation regions. The underlying principle is that individuals with a higher number of KIV-2 repeats will exhibit higher occurrences of KIV-2 kmers. The normalization regions, representing single-copy genomic areas, serve as a calibration factor to account for variations in sequencing depth across individuals.

The KILDA toolkit is organised around *kilda*.*nf*, a Nextflow pipeline. This pipeline can run two subworkflows, (i) *prepare_kmers_DB* to create a list of kmers specific to KIV-2 and the normalisation regions, (ii) *kiv2_counts* to count the number of KIV-2 from FASTQ files. The second component uses a Python script, *kilda*.*py*, to estimate the KIV-2 repeats, which can also be run independently. For ease-of-use KILDA is provided with a list of kmers specific to KIV-2 and the *LPA* gene for the normalisation region, hence running the first subworkflow is optional. The KILDA pipeline is intended to be flexible and the user can decide to run one or both of the subworkflows by setting the corresponding booleans to true or false in the config file.

### Prepare kmers DB

This subworkflow is designed to generate the list of kmers specific to the KIV-2 and normalisation regions (Supplementary Figure 1):

1. Kmers from the reference genome are generated with Jellyfish v2.2.10 [13], based on regions given as bed files.
2. Kmers are filtered based on their occurrences (1 for the normalisation kmers, 6 for the KIV-2 kmers, as there are 6 KIV-2 repeats in the reference genome).
3. The kmers with a count > 0 anywhere else are discarded.
4. Kmers in common between the KIV-2 and normalisation regions are discarded.
5. The kmers are written under the FASTA and tsv formats.

This ensures that we have representative kmers that are specific to the KIV-2 and the normalisation regions. A template configuration file is provided, users can modify the regions of interest, as well as the kmer size.

### KIV2 counts

The second subworkflow from KILDA can infer the number of KIV-2 repeats directly from FASTQ files. It relies on kmers specific to the KIV-2 region and to one or more normalisation regions to estimate the KIV-2 repeats. These lists of kmers can be produced with the subworkflow described previously.

The kmers from both lists are counted with *jellyfish count* and are written to a tab-delimited counts file with *jellyfish dump*. The counts files are then given as input to *kilda*.*py* for estimation of the KIV-2 copy numbers (Supplementary Figure 2).

### kilda.py

The python script reads the counts files and computes the number of KIV-2 repeats by dividing the mean occurrence of the KIV-2 kmers by the mean occurrence of the normalisation kmers. A report for each sample, indicating the number of missing kmers and the occurrence means is printed to the terminal.

If the *--rsids* option was set, *kilda*.*py* will report the number of reference and alternative kmers that were counted for each single nucleotide polymorphism (SNP) given as input by the user. This can help to give more context around the number of KIV-2 repeats, as some SNPs can affect Lp(a) concentration [4]. If the *--plot* option is activated, KILDA generates a visual representation of the distribution of the occurrences of the KIV-2 and normalisation kmers for each sample (Supplementary Figure 3).

*kilda*.*py* can be executed independently or from within *kilda*.*nf*.

### Comparison against DRAGEN-LPA on the 1000 Genomes Project

KILDA was tested against the DRAGEN-LPA caller [12] on the 1000 Genomes Project dataset [14]. The DRAGEN-LPA KIV-2 copy number predictions were obtained from the preprint’s Supplementary Tables. KILDA’s predictions were computed from the 1000G phase 3 FASTQ files, using a list of 21-mers produced with the *prepare_kmers_DB* subworkflow. Additionally, we compared the output of KILDA directly to the “high-confidence” Bionano KIV-2 alleles available from Supplementary Table 1 of the DRAGEN-LPA manuscript [12].

In addition to its primary functionality, KILDA can process a predefined list of variants with corresponding kmers which contain reference and alternative alleles. We thus selected the following variants: *rs10455872* (+5.4 mg/dL), *rs3798220* (+45 mg/dL) and *rs41272114* (−5 mg/dL) with established impact on Lp(a) concentrations [4]. A sample was considered as carrier if at least one alternative kmer was detected in the sample.

### *In-silico* predictions

We created *in-silico* samples with pre-determined KIV-2 copy-numbers to test KILDA’s robustness under different scenarios of sequencing coverage and quality.

First, we created a “diploid” reference, by duplicating GRCh38 and applying dbSNP’s SNPs to one of the copies, using *bcftools consensus*. Then, for each condition (coverage between 2 and 30X, and mean read quality between 12 and 28), we created 40 samples by adding a random number of KIV-2 sequences (between 6 and 30) to each of the haplotypes, resulting in samples with KIV-2 copy numbers between 12 and 60. KIV-2 sequences were randomly selected from the 6 KIV-2 sequences from GRCh38.

### KILDA: identifying KIV-2 repeats from kmers

Finally, we generated whole-genome sequencing data as paired-end reads of 150 bp, using *bbmap randomreads*.*sh* with different options regarding coverage and mean quality. For the coverage (option *– coverage*), a fixed mean quality of 28 was set (the default value of *bbmap randomreads*.*sh*), and we simulated reads with estimated coverages of 2, 5, 10, 20 and 30X. For the mean quality (option *–midq)*, a fixed coverage of 10X was set, and we generated libraries of reads with a mean quality of 12, 14, 16, 18, 20 and 28.

The *kiv2_counts* subworkflow was run on each set of 40 samples, using a list of 31-mers produced with KILDA. The predicted KIV-2 copy numbers were compared to the known number of KIV-2 induced in the samples.

## 3 Results

### Comparison against DRAGEN-LPA on the 1000 Genomes Project

KILDA’s prediction for the 1000 Genomes Project samples were highly correlated to the DRAGEN-LPA predictions with a R^2^ = 0.935 (y = 1.22x - 1) (see Figure 1).

**Figure 1:**
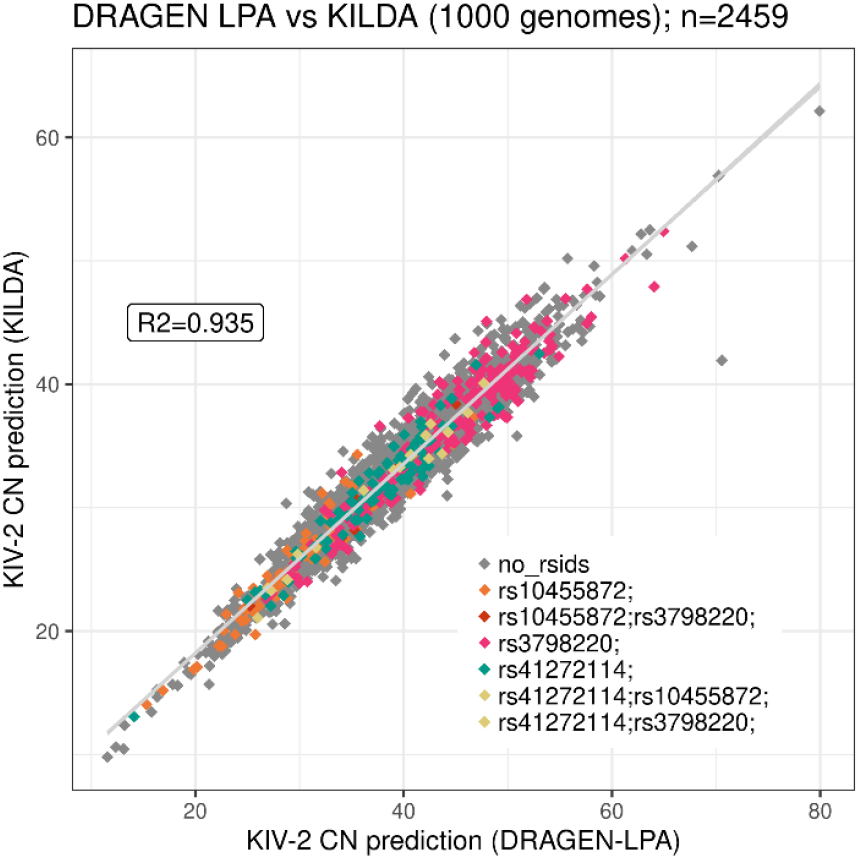
KILDA predictions against DRAGEN-LPA predictions for the number of KIV-2 copies on samples from the 1000 Genomes Projects. For KILDA, 21-mers were used to make the predictions. The samples are colored by the presence of variants in samples as detected by KILDA.

However, there was a factor of 1.22 between the predictions made with KILDA, using 21-mers, and DRAGEN-LPA. Testing with different values of *k* showed that increasing the kmer size resulted in a reduction of this factor (Supplementary Figure 4), at the cost of more computational resources. If reads are of high quality and computational resources are available, we recommend using higher values of *k* when using KILDA. KILDA’s prediction were also correlated to the “highconfidence” Bionano KIV-2 set from DRAGEN-LPA Supplementary Table 1 (R^2^ = 0.966, y = 1.22x – 1.2). The comparison of KILDA’s prediction against the optically mapped samples is shown in Supplementary Figure 5.

Three variants were given as input to KILDA to give more context when interpreting the number of KIV-2 in relation to Lp(a): *rs10455872, rs3798220* and *rs41272114*, which were found in 92, 224 and 105 samples respectively. These samples overlapped well with the carriers of these SNPs in the 1000 Genomes Project VCF file, detailed counts are available in Supplementary Table 1.

The mean number of KIV-2 repeats was different between the carriers and non-carriers for *rs10455872* (26.22 *vs*. 32.95, p-value < 2.2e-16) and *rs3798220* (36.45 *vs*. 32.32, p-value = 1.47e-11). Concerning *rs41272114*, the mean number of repeats was similar between carriers and non-carriers (31.55 *vs*. 32.75, p-value = 0.02). Thus, as previously described [4] *rs10455872* is partially tagging smaller isoforms. However, *rs3798220*, a Lp(a)-increasing variant, previously thought to be associated with lower isoforms [4], is present in samples with higher number of genetic KIV-2, which could be wrongly interpreted as having lower Lp(a) and lower risk of CVD, demonstrating the importance of including the presence of Lp(a)-affecting variants when interpreting the number of KIV-2 copies. The discordance between our results and the literature can be explained when stratifying the results by ancestry. *rs3798220* is indeed present in smaller isoforms in Europeans (28.17 *vs* 31.56) but associated with longer isoforms in Americans (36.63 *vs* 31.78) which are the main carriers of this SNP (n=115). In addition, this SNP is rare in Africans who have smaller isoforms (Supplementary Figure 6). Highlighting the importance of directly estimating KIV-2 repeats instead of uniquely relying on rsids when studying diverse populations.

The pipeline was run on a Debian 11 server, with 5 CPUs allocated to each sample. Each sample was processed in 18 minutes, with a peak memory usage of 830 Mb. The counting of the kmers with *jellyfish count* being the most time-consuming (Supplementary Table 2).

### *In-silico* predictions

The R^2^ between the predicted KIV-2 copy numbers and the copy numbers included in the simulated samples remained higher than 0.99 for coverages between 5X and 30X (Supplementary Figure 7). For 2X, the lowest tested coverage, the R^2^ value was 0.97, demonstrating KILDA’s accuracy even at very low coverages.

Regarding the reads mean quality, the R^2^ between the predicted and simulated KIV-2 copy numbers was > 0.99 for mean sequencing qualities between 18 and 28 (Supplementary Figure 8). KILDA’s accuracy remained high for libraries with mean qualities of 16 and 14 (R^2^ = 0.98 and 0.95 respectively), showing KILDA’s robustness to sequencing libraries of lower quality. KILDA’s predictions started to falter when simulating a library with a mean quality of 12, but realistically, modern sequencing libraries should reach higher mean qualities.

## 4 Conclusion

Here, we demonstrated the performance of KILDA on a real whole genome sequencing dataset: the 1000 Genomes Project, and against simulated datasets with known number of KIV-2 repeats. Future work should focus on reducing the factor of prediction observed against DRAGEN-LPA, and on including haplotyping of the KIV-2 copy numbers, as it can influence Lp(a). To further validate KILDA’s utility and applicability in clinical settings, it is imperative to evaluate its performance on whole exome sequencing and targeted panel datasets. This additional testing would help determine whether KILDA can be effectively integrated into routine diagnostic practices. It would therefore provide a free tool complementary to that of variant calling within unmappable regions [15] using short read sequencing, instead of long read [16], which is currently not commonly used in clinical practice.

KILDA is an open-source Nextflow pipeline, which can estimate the number of KIV-2 repeats along with the presence of Lp(a)-affecting variants. One of the main advantages of KILDA is its capacity to estimate KIV-2 numbers based on the *LPA* sequence alone and so does not need normalisation regions across the genome which are often not captured in clinical practice. KILDA is provided with an Apptainer image and pre-determined lists of KIV-2 and normalisation kmers for ease-ofuse.

## Acknowledgement

We would like to thank Pierre Lindenbaum for his feedback during KILDA’s development.

