## Supplementary Materials for "KILDA: identifying KIV-2 repeats from kmers"

“KILDA: Detecting KIV-2 copy numbers from sequencing data”

Supplementary Materials

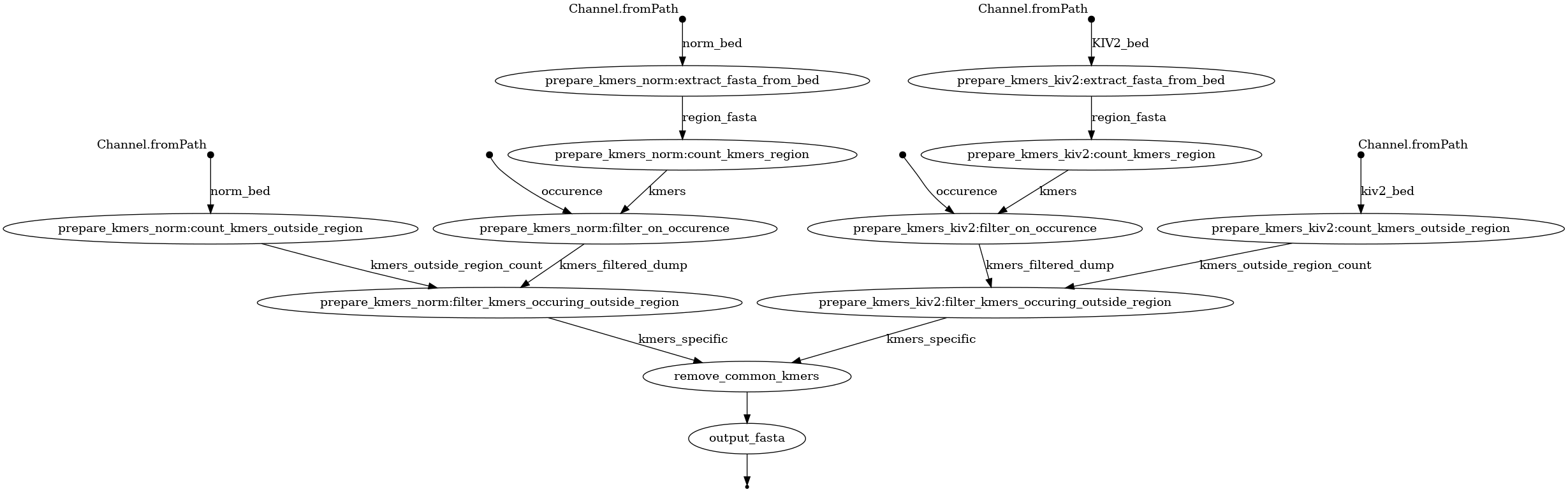

Supplementary Figure 1: “kiv2_create_kmers_DB.nf” workflow. The KIV-2 and normalisation regions are extracted from the reference genome using coordinates given as bed files. Then the kmers are counted on the region and are filtered on their occurrence (only keeping kmers with occurrence 6 for KIV-2 and occurrence 1 for the normalisation region). Remaining kmers with an occurrence > 0 on the regions outside the regions of interest are removed to only keep kmers specific to KIV-2 and the normalisation region. Finally, any kmer found in common between the KIV-2 list and the normalisation list is removed. The final list of kmers is written as FASTA and tsv files.

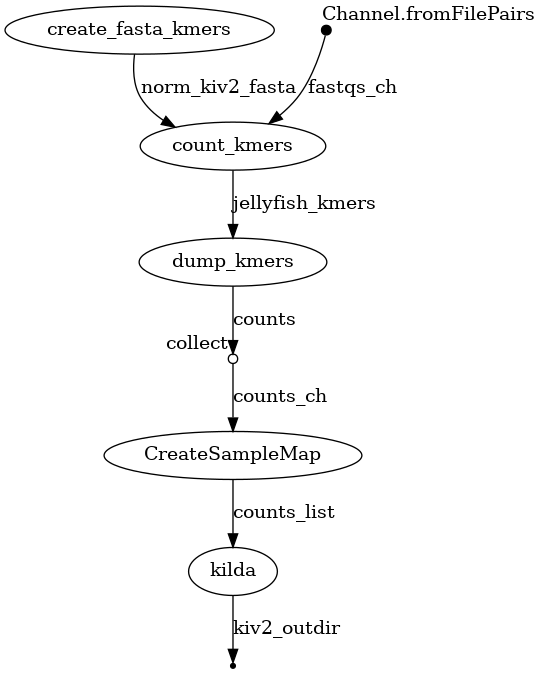

Supplementary Figure 2: First, kmers from KIV-2 and the normalisation region are written to a FASTA file, which is used as argument to “jellyfish count” to limit counting to the kmers of interest. The counts are then dumped to a tsv file and the results for all the samples are collected and written to a “sample map” linking the sample identifiers to their corresponding dump file. The sample map is given to “kilda.py” for estimation of the KIV-2 copy number.

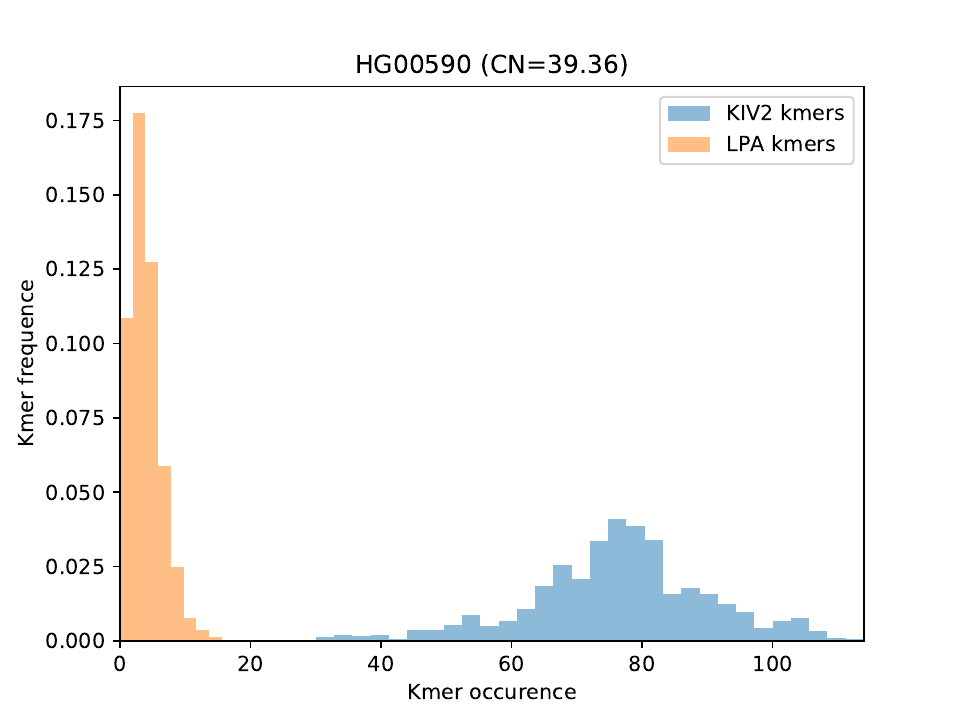

Supplementary Figure 3: Example of plot generated by KILDA for sample “HG00590” from the 1000 Genomes project. This plot shows the distribution of the occurrences of the KIV-2 kmers (in blue) and the normalisation kmers (in yellow). KILDA will determinate the number of KIV-2 repeats (here 39.36) based on the ratio between the mean occurrence of the KIV-2 kmers and the normalisation kmers (here the LPA gene).

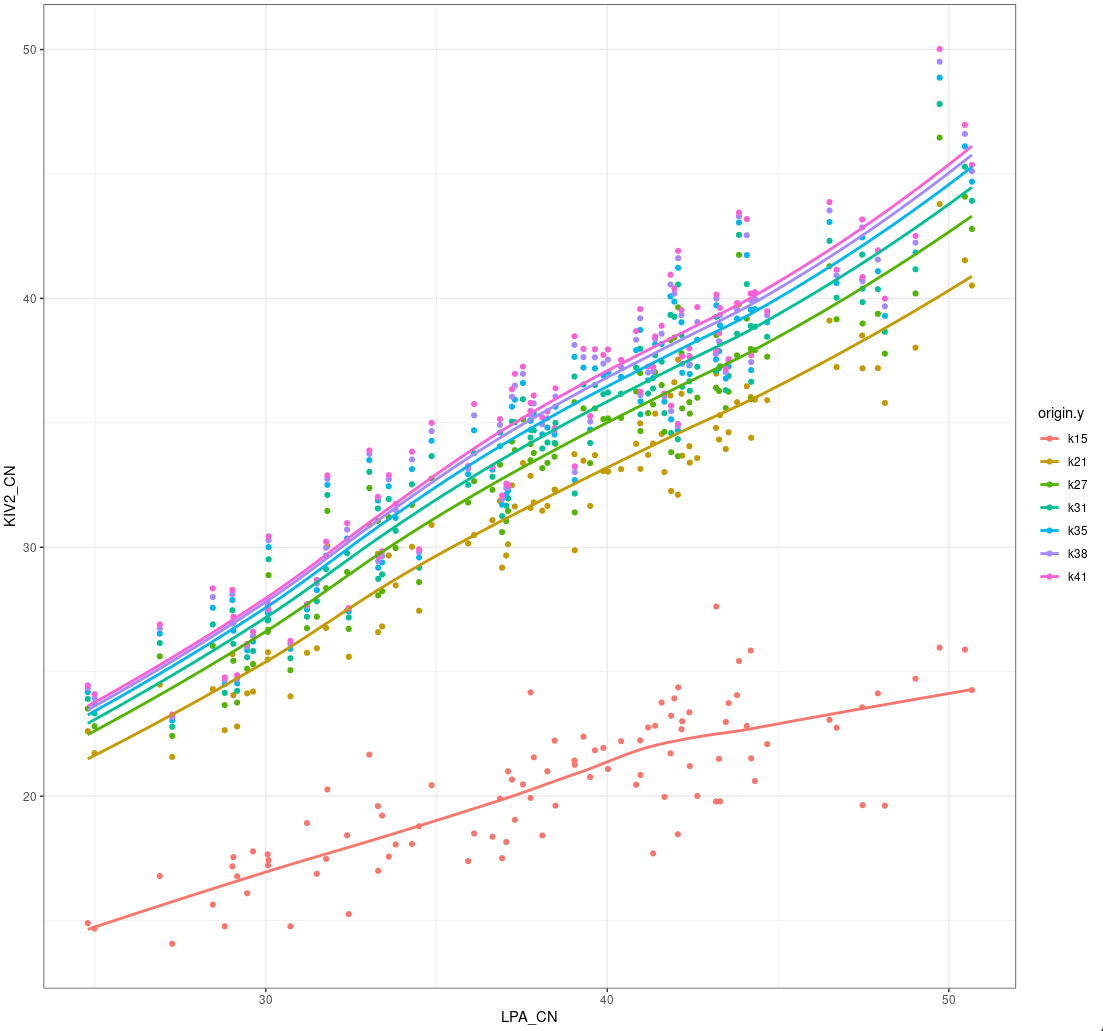

Supplementary Figure 4: Impact of the kmer size on KILDA's predictions. For k=15 the kmers are small and not specific enough and KILDA’s predictions are not accurate. From k=21 to k=41 predictions remain proportionate but with an increase in the copy numbers with an increase in k.

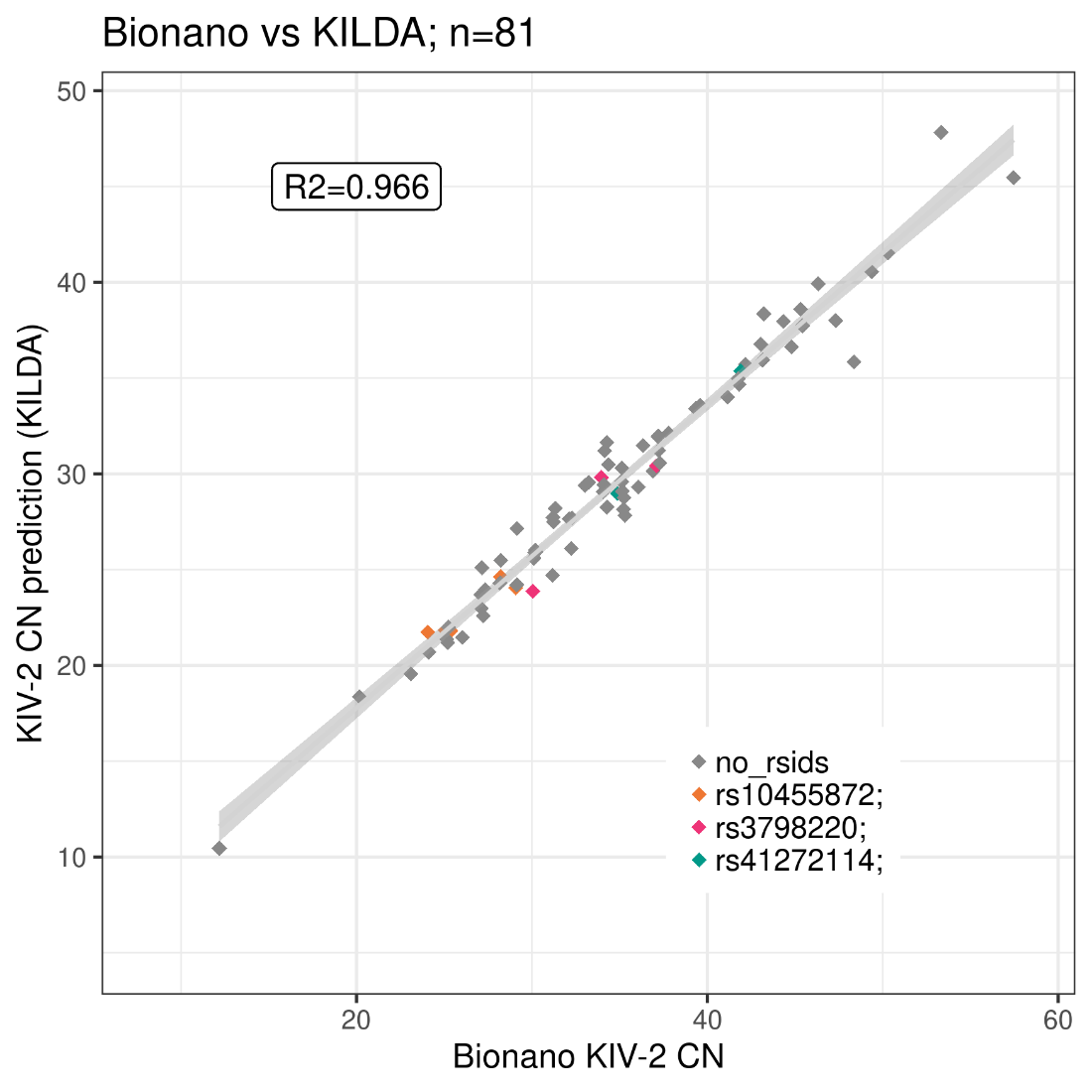

Supplementary Figure 5: KILDA’s KIV-2 copy number predictions against the “high-confidence” Bionano optical maps of the LPA region. For Bionano, the number of KIV-2 alleles was retrieved from Supplementary Table 1 of the DRAGEN-LPA manuscript. For KILDA, 21-mers were used to make the predictions. The samples are colored by the presence of variants in samples, as detected by KILDA.

Supplementary Table 1: Overlap between the SNPs carriers detected with KILDA against the carriers present in the 1000 Genomes Project phase 3 VCF file. **A.** overlap for rs10455872 **B.** overlap for rs3798220 **C.** overlap for rs41272114

**A.**

| **rs10455872** | *VCF carriers* | *VCF non-carriers* |
| --- | --- | --- |
| *KILDA carriers* | 85 | 7 |
| *KILDA non-carriers* | 19 | 2332 |

**B*.***

| **rs3798220** | *VCF carriers* | *VCF non-carriers* |
| --- | --- | --- |
| *KILDA carriers* | 207 | 15 |
| *KILDA non-carriers* | 24 | 2197 |

**C*.***

| **rs41272114** | *VCF carriers* | *VCF non-carriers* |
| --- | --- | --- |
| *KILDA carriers* | 97 | 6 |
| *KILDA non-carriers* | 13 | 2327 |

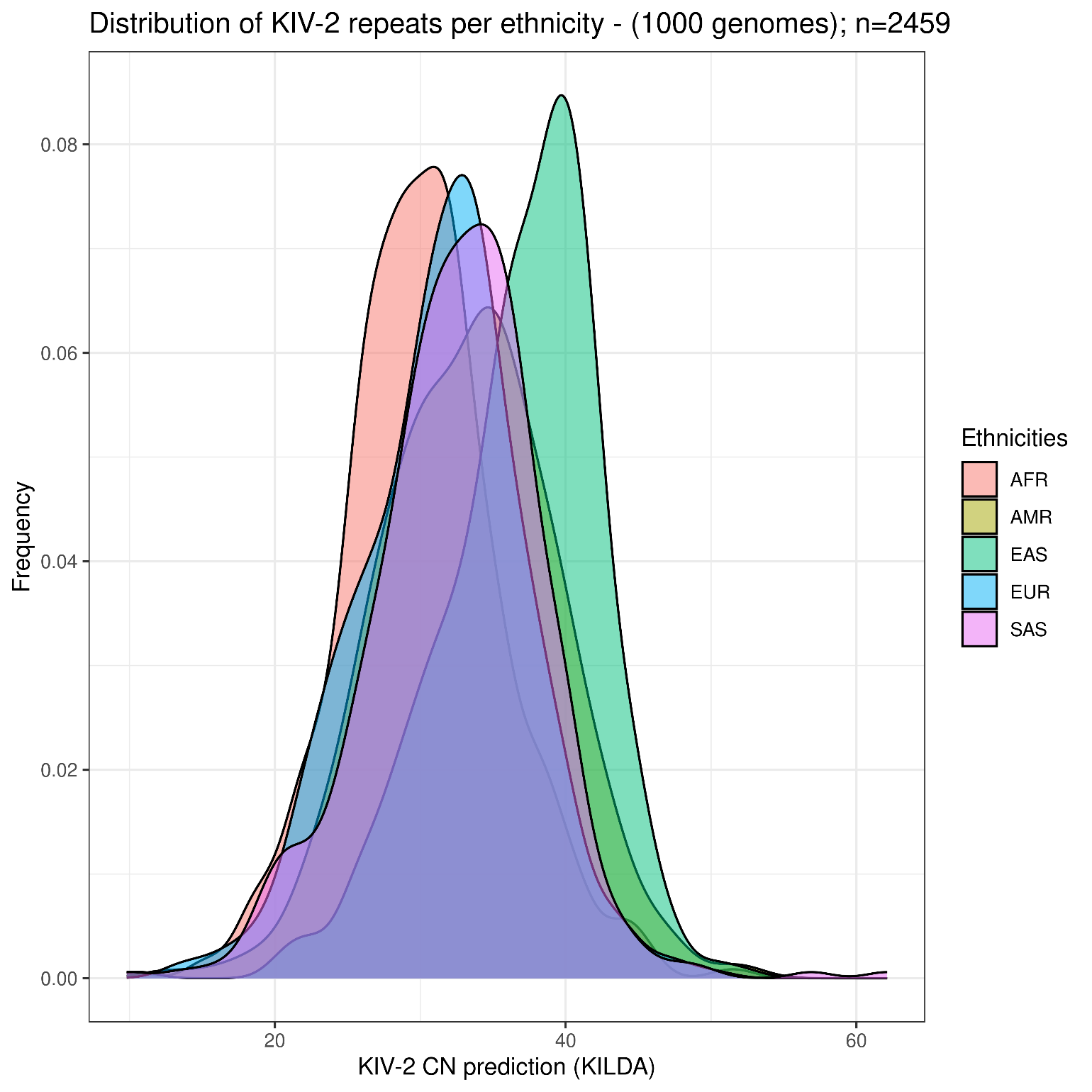

Supplementary Figure 6: Distribution of KILDA's KIV-2 copy number predictions per population, as represented in the 1000 Genomes project (AFR = African, AMR = American, EAS = East Asian, EUR = European, SAS = South Asian).

Supplementary Table 2: Nextflow's statistics upon running KILDA on the 2,239 samples from the 1000 genomes project. For each step the mean memory usage and job duration is detailed. The steps correspond to the Nextflow pipeline shown in Supplementary Figure 2.

| **Step (number of jobs)** | **Mean Memory Usage (M)** | **Mean Job Duration (minutes)** |
| --- | --- | --- |
| create_fasta_kmers (1) | 0 | 0 |
| count_kmers (2239) | 830 | 14 |
| dump_kmers (2239) | 2 | 0 |
| CreateSampleMap (1) | 0 | 0 |
| kilda.py (1) | 195 | 19 |

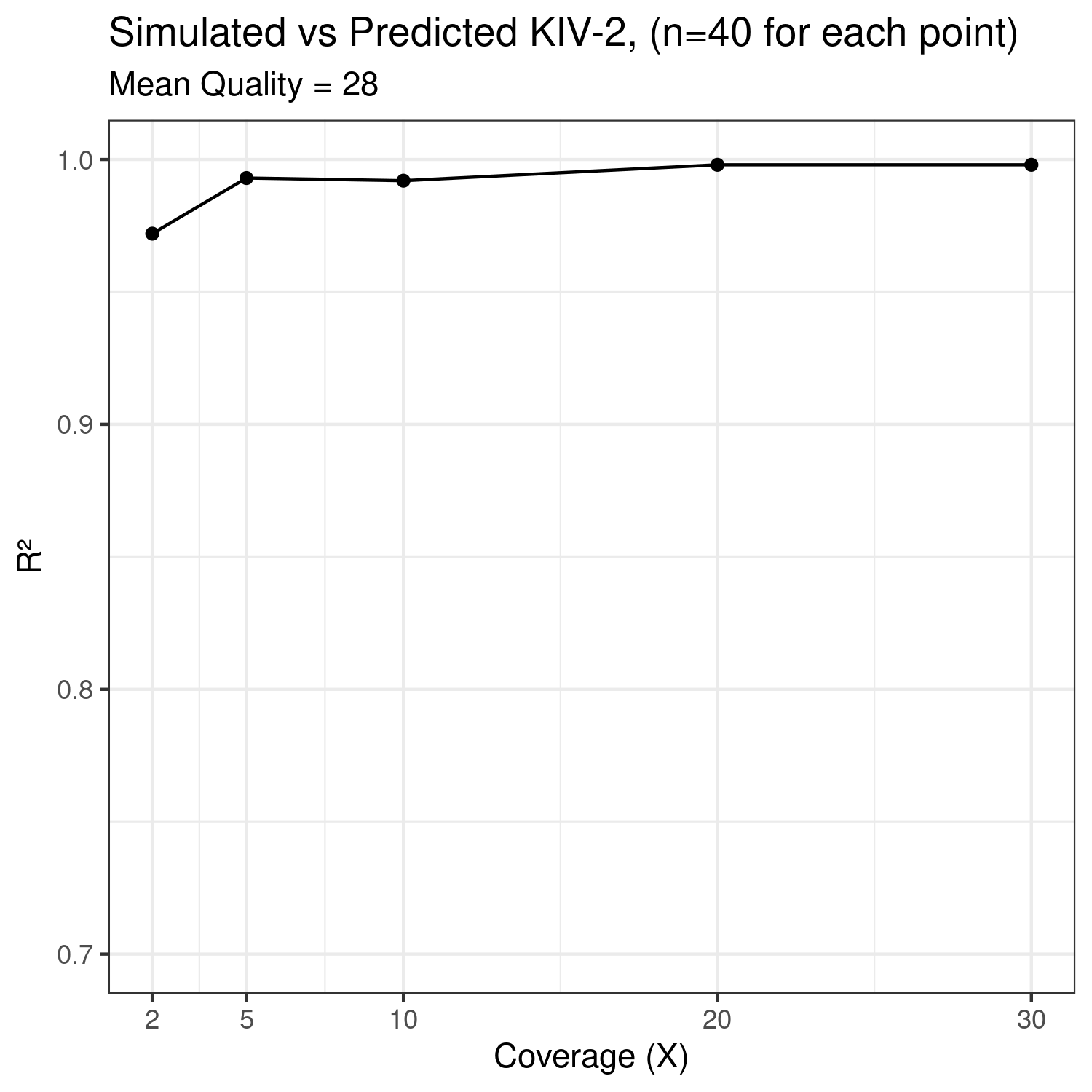

Supplementary Figure 7: Computed R^2^ between the number of KIV-2 detected with KILDA against the number of KIV-2 inserted in-silico into each sample (n=40) for simulated sequencing libraries with different coverage values. KILDA maintains a high accuracy to detect the correct number of KIV-2 repeats even at low coverages (R^2^=0.97 at 2X). The mean read quality was set to the default value (28) of “bbmap randomreads.sh“.

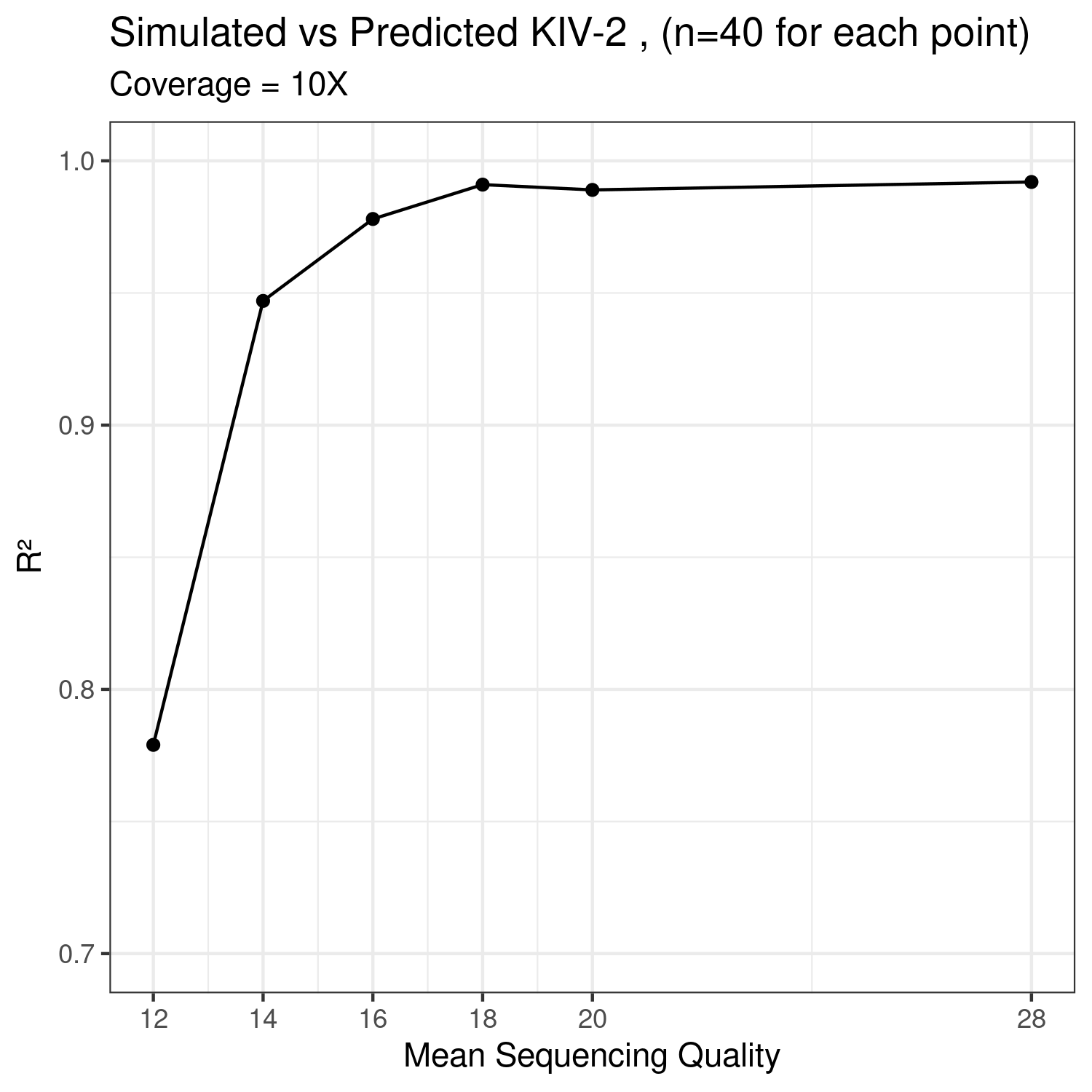

Supplementary Figure 8: Computed R^2^ between the number of KIV-2 detected with KILDA against the number of KIV-2 inserted in-silico into each sample (n=40) for simulated sequencing libraries with different mean quality values. KILDA maintains a high accuracy for standard mean read quality (>20). A fixed coverage value of 10X was set for all comparisons.
